# Analysis of SFTS pathogenesis based on single-cell RNA sequencing of monocytes

**DOI:** 10.1101/2025.02.26.640382

**Authors:** Yin Zhang, Mengtao Gong, Qianqian Yang, Yangyang Jin, Yuanhong Xu

## Abstract

**Backgroud:** Severe fever with thrombocytopenia syndrome (SFTS) is a newly identified tick-borne viral disease with a high mortality rate. Monocytes are known to play an important role in the pathogenesis of SFTS, but the distinct subpopulations of monocytes associated with the disease progression and their roles in inflammatory responses remain unclear. In this study, we used single-cell RNA sequencing (scRNA-seq) to explore the specific monocyte subpopulations correlated with the progression of SFTS and their involvement in inflammatory processes.

**Methods:** scRNA-seq analysis was conducted on isolated monocytes from seven SFTS patients, validated by cell identification through flow cytometry. Comparative analyses were performed using three normal monocyte samples from public databases. Bioinformatic tools including Cell Ranger, Seurat, Monocle 2, GSEA, and SCENIC were utilized to identify distinct cell populations and transcriptional patterns.

**Results:** Our results indicate that severe SFTS patients in the ICU exhibited a reduced proportion of circulating HLA-DR^+^CD14^+^ monocytes compared to non-severe hospitalized SFTS patients, indicating a dysfunctional immune response. Increased numbers of CD163^+^CD14^+^ monocytes in acute SFTS patients correlated with disease severity. CD163^+^CD14^+^ monocytes were identified as potential target cells for SFTSV infection. SFTSV infection could drive monocyte differentiation into the M2 phenotype, promoting virus persistence and disease advancement.

**Conclusion:** Our studies in SFTS patients have shown that monocytes are target cells for SFTSV. Notably, an expansion of CD163^+^CD14^+^ monocytes in SFTS has been shown to be associated with severe clinical disease. In addition, activated IL-6/JAK/STAT3 signaling in SFTS patients induced CD163^+^CD14^+^ monocyte expansion, which was recovered by tofacitinib treatment.

## Introduction

Severe fever with thrombocytopenia syndrome (SFTS) is caused by the SFTS virus (SFTSV), transmitted through ticks and first identified in rural central China in 2010 [1]. The mortality rate of SFTS ranges from 12% to 30% based on previous cases in China, South Korea, and Japan. Patients typically exhibit non-specific symptoms such as fever, thrombocytopenia, leukopenia, and systemic manifestations including hemorrhagic symptoms. Severe patients worsen rapidly within a week post-symptom onset, often died of multi-organ failure.

During the acute phase of SFTS infection, SFTSV triggers an immune response in monocytes, leading to the dysregulated release of immune factors like tumor necrosis factor-α (TNF-α), interleukin-1β (IL-1β), and IL-6, contributing to tissue damage [6]. Monocytes, important components of the innate immune system, recognize pathogens via Toll-like receptors (TLRs), triggering the secretion of cytokines, chemokines, and signaling molecules. They can differentiate into macrophages or myeloid dendritic cells (mDCs) to initiate immune responses. Early in severe SFTSV infection, researchers noted substantial monocyte apoptosis and necrosis, indicating compromised immunity post-infection [7]. Additionally, infected monocytes showed reduced response to LPS stimulation compared to patients in recovery or healthy controls, suggesting weakened immune function [8]. Moreover, previous studies have shown blood monocytes as targets for various viruses like dengue and Zika, with viral infections triggering the transformation of monocytes into dendritic cells [9–11].

While monocytes play critical roles in SFTS progression, there’s limited understanding regarding the diverse functions of distinct monocyte subpopulations in SFTS. Their exact functions, gene expression patterns, and activation statuses remain unclear. Advances in single-cell RNA sequencing have enabled more accurate comparative analysis of mononuclear cells. Therefore, our aim was to provide a detailed characterization of monocyte subpopulations and their specific gene expression profiles in SFTS, with the goal of identifying the genes and markers that drive SFTS progression.

## Methods

### Patient cohort, sampling and data collection

We recruited adult patients with laboratory-confirmed SFTSV infection who were admitted to the hospital between April 28 and August 29, 2024. The SFTSV infection was confirmed using quantitative reverse transcription polymerase chain reaction (qRT-PCR) based on established criteria [12]. Healthy individuals of similar age, admitted for health assessments, and confirmed negative for SFTSV infection, were included as controls during the study period.

All study procedures were in accordance with the Declaration of Helsinki and approved by the Ethics Committee of the First Affiliated Hospital of Anhui Medical University (Authorization number: PJ 2024-09-51, Hefei, China).

The SFTS was defined as a critical illness when; the patient was admitted to the intensive care unit (ICU), had mechanical ventilation, or died. Blood samples from all patients were collected at the earliest possible timepoint after admission, as per study protocol. Separation of plasma and PBMCs from EDTA tubes was performed using a lymphocyte separation medium (TBD Science).

### SFTS viral load assay

As described previously [12], we extracted total RNA from the whole blood samples of patients using a viral RNA kit (DaAn Gene, Guangzhou, China) and quantified the SFTSV RNA copies with a validated real-time PCR kit (DaAn Gene, Guangzhou, China) according to the manufacturer’s protocols.

### Laboratory index detection

The peripheral blood of 2 ml patients was placed in the EDTA anticoagulant tube. Subsequently, white blood cells, neutrophils, lymphocytes, monocytes, platelets, and hemoglobin levels were analyzed using the XN9000 fully automatic analyzer (Sysmex, Japan) along with the respective accompanying reagents.

### PBMC isolation

Peripheral blood mononuclear cells (PBMCs) were isolated using lymphocyte separation medium (TBD Science) via density gradient centrifugation and subsequently washed with Ca/Mg-free PBS (Procell). The PBMCs underwent treatment with RBC Lysis Buffer (Solarbio) to eliminate red blood cells, followed by washing with Ca/Mg-free PBS supplemented with 2% fetal bovine serum (Gibco). A portion was used for cell counting via the Countess II Cell Counter (Thermofisher). The viability of all freshly isolated PBMCs was over 90%.

### Flow cytometry

Peripheral blood samples were processed within 2 hours for the analysis of lymphocyte/monocyte subsets using flow cytometry. PBMCs were pre-treated with human Fc block (1:100, BD Biosciences) for 10 minutes at 4°C. Each sample received an appropriate antibody cocktail for staining surface markers, with a staining duration of 30 minutes at 4°C. For identifying lymphocyte subsets among PBMCs, antibodies like CD3-PE/Cyanine7, CD8-APC-H7, CD19-APC, and CD56-BV421 were used. In profiling monocyte subsets, antibodies such as CD14-BV605, CD16-PE/Cyanine7, HLA-DR-PerCP-Cy5.5, CD163-AF647, and CD163-FITC (BD Pharmingen) were utilized. All fluorescent antibodies were sourced from BioLegend, unless specified otherwise.

Intracellular cytokine staining involved incubating PBMCs with a specific solution at 37°C in a 5% CO2 incubator. After staining with surface antibodies, cells were fixed, permeabilized, and stained with antibodies targeting various intracellular cytokines. For intracellular phosphoprotein evaluation, cells were fixed, permeabilized, and stained with Phospho-STAT3 antibodies. Analysis was carried out using a BD FACS Canto II and FlowJo Software

### Monocyte isolation and purification

Monocytes were extracted from PBMCs of healthy donors using the Pan-monocyte Isolation Kit (Miltenyi) following the manufacturer’s guidelines. Post-isolation, the purity of the monocyte populations, evaluated using CD14 and CD16 markers, exceeded 95%.

### Single-cell RNA library preparation and sequencing

The sorted monocyte suspension was loaded onto a Chromium Single Cell Chip (Chromium Single Cell B Chip Kit, 10x Genomics, PN-1000074) using a Chromium Single Cell 3’ GEM, Library & Gel Bead Kit v3 (10x Genomics, PN-1000077) as per the manufacturer’s instructions. Encapsulation with barcoded Gel Beads aimed for around 6000 individual cells per sample. Subsequently, RNA release occurred through reverse-transcription within single-cell gel beads in the emulsion (GEMS). Following cDNA synthesis and amplification via reverse-transcription on a T100 PCR Thermal Cycler (Bio Rad), sequencing took place on an Illumina Novaseq 6000.

### Data quality control

Raw BCL files were processed into FASTQ files, aligned, and quantified using the 10X Genomics Cell Ranger software (version 3.1.0). Removing reads with low-quality barcodes and UMIs was followed by mapping to the reference genome. Reads uniquely mapped to the transcriptome and intersecting an exon at least 50% were considered for UMI counting.

### Cell clustering and annotation

Based on the top 25 principal components, cell clustering utilized the graph-based algorithm in the FindClusters function with a resolution of 0.4. Visualization was enhanced using UMAP. For the sorted monocyte scRNA-seq data, cell clusters were named based on the top DEGs. Differentially expressed genes within a specific cluster were identified by comparing them with other clusters using the Wilcoxon test in the FindAllMarkers function.

### Differentially expressed genes analysis

Seurat facilitated the analysis of differentially expressed genes. New idents were assigned for group_cluster or group_cell type for further analysis [13]. A hurdle model in MAST was employed to identify differentially expressed genes within a group in one cluster based on specific criteria.

### Pathway enrichment analysis

Enriched pathways were evaluated through hypergeometric testing in the Gene Ontology (GO) and Kyoto Encyclopedia of Genes and Genomes (KEGG) databases. Using the compareCluster function in the clusterProfiler package, significantly enriched pathways were determined with a Benjamini–Hochberg corrected p-value threshold [13].

### Gene set variation analysis (GSVA)

Gene Set Variation Analysis (GSVA) was conducted using gene sets from MsigDB to identify enriched pathways and cellular processes across various clusters [14]. This analysis, carried out with the GSVA R package, relied on the log-transformed expression matrix averaged per cluster [15].

### Pseudo-time analysis

#### Constructing single cell trajectories

Analysis of single-cell trajectories involved utilizing the matrix of cells and gene expressions with Monocle [16] (Version 2.10.1). Monocle compressed the multidimensional space into two dimensions and arranged the cells in a specific order (sigma = 0.001, lambda = NULL, param.gamma = 10, tol = 0.001) [17]. This ordering allowed visualizing the trajectory in reduced dimensional space, exhibiting a tree-like structure comprising tips and branches.

#### Differential expression analysis

Monocle was employed to identify genes showing differential expression between cell groups and evaluate the statistical significance of these changes. We pinpointed key genes associated with developmental processes and differentiation with an FDR threshold <1e-5. Genes displaying similar expression patterns were grouped together under the assumption that they might share common biological functions and regulators.

#### Analyzing branches in single-cell trajectories

Single-cell trajectories often contain branches, representing alternative gene expression programs within cells. These branches manifest during development as cells decide their fate: one lineage follows a particular path, while the other lineage takes a different course. Monocle introduced BEAM to examine branch-dependent gene expression by framing the issue as a contrast between two negative binomial GLMs [18].

#### Regulons analysis

The SCENIC [19] tool was utilized to pinpoint differentially activated transcription factors (TFs) in CD163^+^ Monocytes between individuals with SFTS and healthy controls (HC). Initially, gene co-expression networks were identified through GENIE3 [20]. Subsequently, each module was refined based on regulatory motifs near transcription start sites using RcisTarget. Specifically, networks were preserved if the TF-binding motif was enriched among its targets, while genes lacking direct TF-binding motifs were eliminated. The preserved networks were denoted as regulons. Finally, the activity of each regulon for individual cells was assessed through AUC scores employing the AUCell R package. Visualization of gene regulatory network (GRN) plots for all regulons was accomplished using the Cytoscape software [21].

#### Cell culture and stimulation conditions

In the drug treatment assay, IL-6 (25 ng/ml, PeproTech) with or without tofacitinib (10nM, Selleck) was added for stimulation. Cells were harvested for flow cytometry analyses of CD163^+^CD14^+^ monocytes expression after 6 h.

#### Cytokine analysis

Serum samples from both patients and healthy individuals were analyzed using a multiplex microsphere flow immunofluorescence assay with a 12-cytokine detection kit obtained from Tianjin Quanto Biotechnology (Tianjin, China). All procedures strictly followed the manufacturer’ s instructions. Initially, 25 μ L of serum sample was mixed with 75 μ L of sample solution containing 25 μ L of assay buffer, 25 μ L of premixed magnetic beads, and 25 μ L of specific cytokine detection antibodies (targeting IL-1β, IL-6, IL-12, IL-17, IL-8, IL-5, IL-2, IL-4, IL-10, TNF-α, IFN-α, and IFN-γ).

After thorough mixing on a plate shaker at approximately 400-500 rpm for 2 hours at 25℃, each test tube received an additional 25μ L of SA-PE and was further agitated (around 400-500 rpm) for 30 minutes at 25℃. Subsequently, 500μL of 1x wash buffer was added to all tubes, followed by centrifugation for 5 minutes at 300-500 g to remove the supernatant. The next step involved adding 100 µL of 1x wash buffer to resuspend the beads by vortexing for 30 seconds. Finally, the samples were analyzed using a Beckman DxFLEX flow cytometer.

### Statistical analysis

Statistical analyses were carried out using GraphPad Prism (GraphPad Software, USA). For comparing two groups, independent-sample t-tests or paired t-tests were applied to normally distributed variables, while Wilcoxon rank-sum tests were utilized for non-normally distributed variables. When assessing more than two groups, one-way analysis of variance (ANOVA) with the Tukey-Kramer post hoc test was employed for data exhibiting normal distribution and homogeneity of variance. A two-tailed p-value less than 0.05 was considered indicative of statistical significance, where *, **, ***, and **** denote p<0.05, p<0.01, p<0.001, and p<0.0001, respectively.

## Results

### Correlation between viremia and inflammatory cytokines

The pathogenesis of severe SFTSV infection usually associates with enhanced viral replication, leading to elevated viremia and a cytokine storm [22]. After confirming acute SFTSV infection in 63 patients, levels of viral load and cytokines were measured in serum samples. Compared with healthy controls, the expression levels of IL-1β, IL-6, IL-10, IFN-α and IFN-γ in SFTSV-infected patients were significantly increased (Fig. 1a-e). IL-6 levels were particularly increased in SFTS patients, acting as a key medium within the acute inflammatory response and the characteristic cytokine storm features. Conversely, levels of IL-2, IL-4, IL-5, IL-8, IL-12P70, IL-17, and TNF-α did not show significant differences between SFTS patients and healthy controls.

**Fig. 1.**
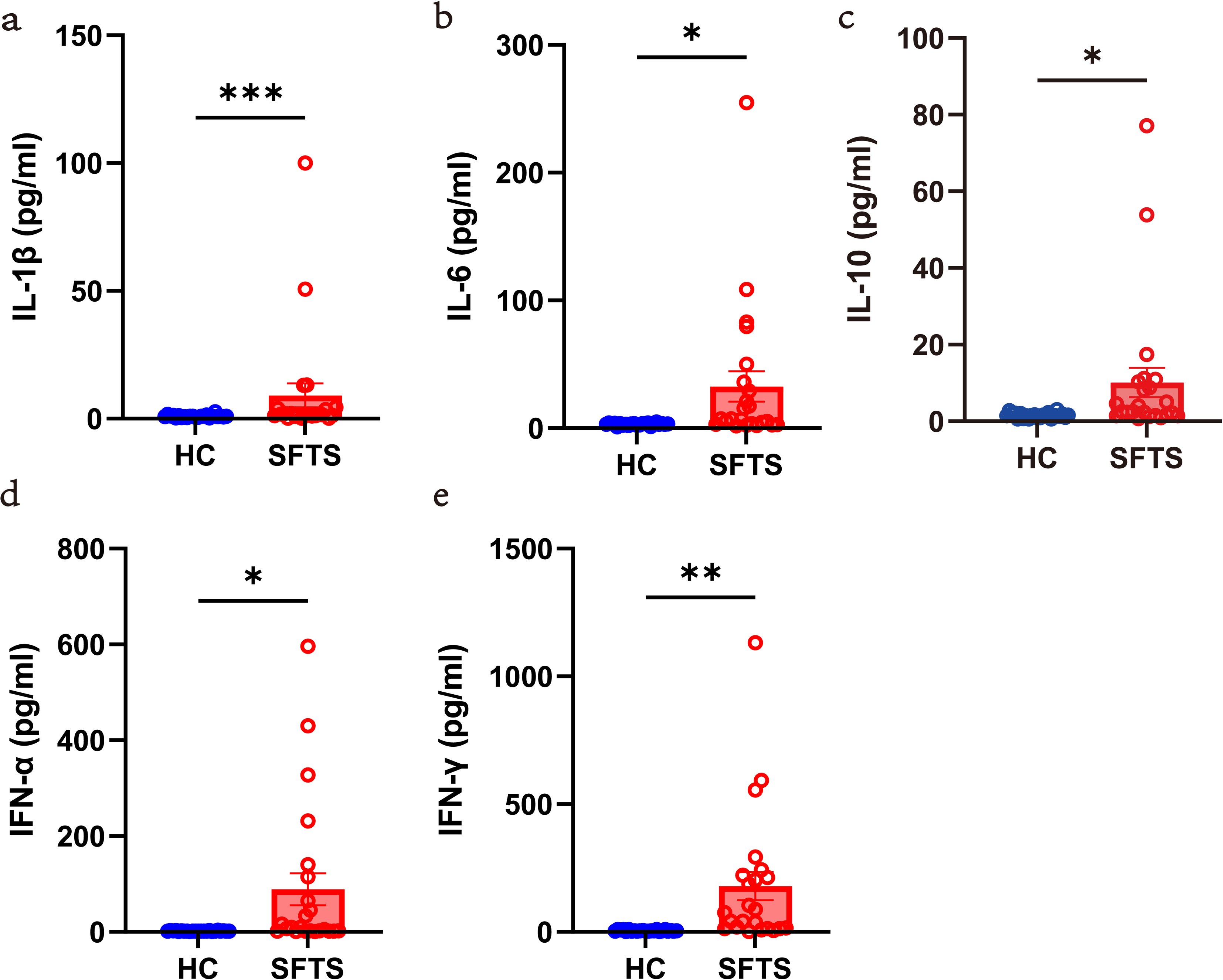
SFTS patients got cytokine storms. Comparison of cytokine levels of (a) IL-1β, (b) IL-6, (c) IL-10, (d) IFN-α and (e) IFN-γ between the HC group and SFTS patients. Data are presented as the mean±SEM. Either Student’s t-test or the Mann-Whitney U test was performed, according to the data distribution. \**p*< 0.05, \*\**p*≤ 0.01, \*\*\*\**p*≤ 0.0001.

To determine whether cytokine responses early during SFTSV infection correlated with an increase in plasma viral load (PVL), we performed correlation coefficient analyses between PVL and each significantly increased cytokine during acute SFTSV infection. Interestingly, a positive correlation emerged between IL-6 and PVL (r = 0.37, p < 0.01, Fig. 2a), IL-10 and PVL (r = 0.51, p < 0.0001, Fig. 2b), and IFN-γ and PVL (r = 0.46, p < 0.0001, Fig. 2c). These findings suggest a potential interplay where elevated PVL may influence cytokine upregulation or vice versa, influencing viral replication. However, no significant correlation was observed between SFTSV load and IFN-α levels (r = 0.03, p = 0.85, Fig. 2d).

**Fig. 2.**
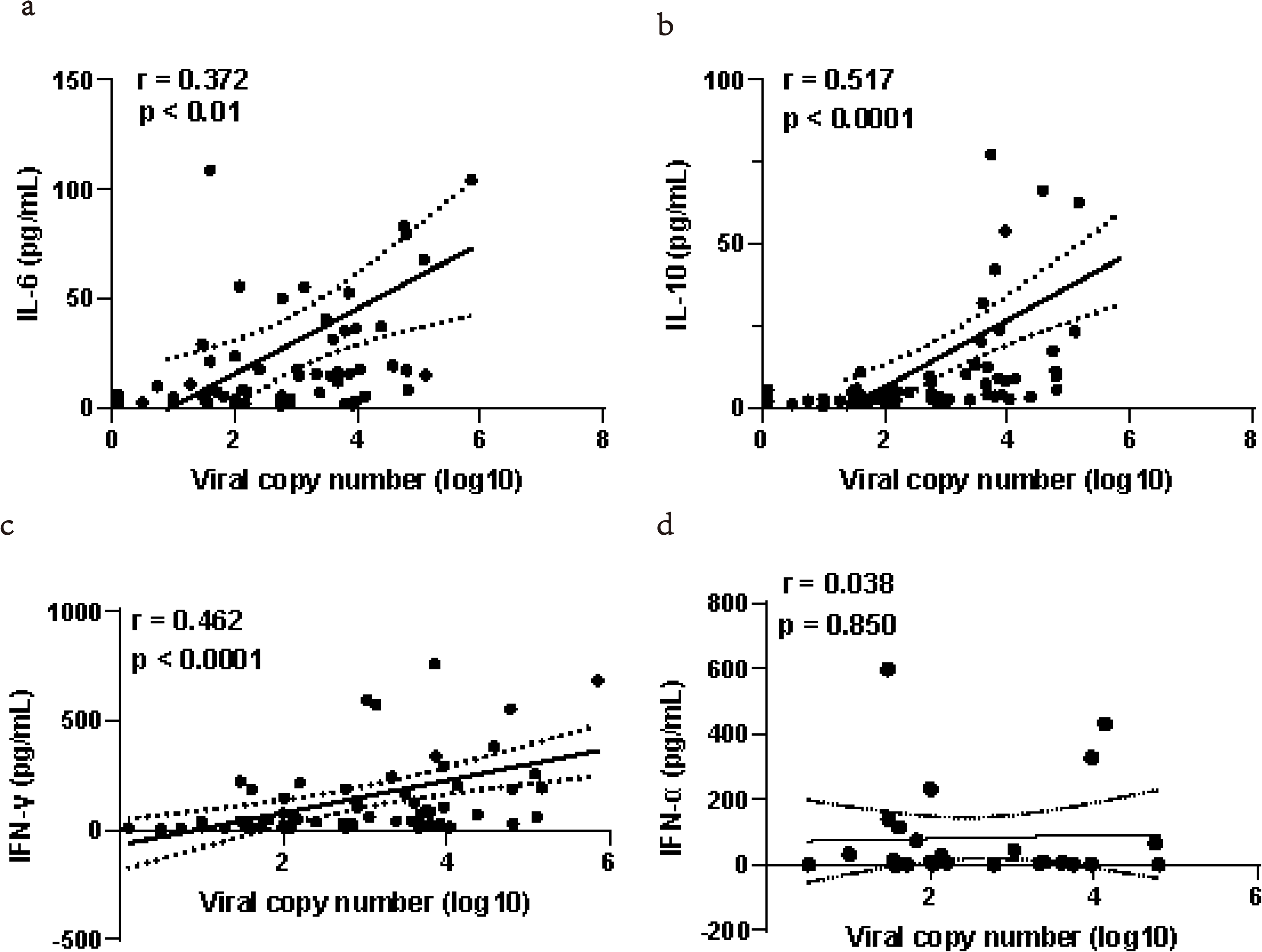
Correlation of cytokine with plasma viral load (PVL). Spearman’s rank correlation coefficient of determination between IL-6 and PVL (a), IL-10 and PVL (b), and IFN-γ (c), and IFN-α (d) and PVL is shown for all SFTSV-infected patients. A significant positive correlation was detected with the expression of IL-6, IL-10, and IFN-γ with PVL during SFTSV infection. R2 and the exact two-sided p value calculated by a Pearson test are shown for correlation analysis.

### PBMCs in peripheral blood

In our study aiming to comprehensively analyze the lymphocyte composition of PBMCs in SFTS patients, we utilized flow cytometry to examine subsets’ absolute numbers and percentages. Notably, lymphocyte counts decreased in PBMCs of SFTS patients, a trend that persisted post-discharge (Fig. 3a). The ratio and cell count of CD3^+^ T cells in SFTS patients exhibited a decline, which subsequently recovered to normal levels during convalescence (Fig. 3c,d). While the T-cell population ratio remained stable in SFTS patients (Fig. 3e,f), the counts of CD4^+^ T cells, CD8^+^ T cells, and the CD4/CD8 ratio decreased (Fig. 3g,h,i). Regarding NK cells, there was no significant difference in the ratios among the healthy control group, SFTS patients, and convalescent individuals (Fig. 3j), but the numbers of NK cells notably dropped during the acute phase (Fig. 3k). Moreover, similar to NK cells, B cells exhibited a comparable trend following SFTSV infection (Fig. 3l,m).

**Fig. 3.**
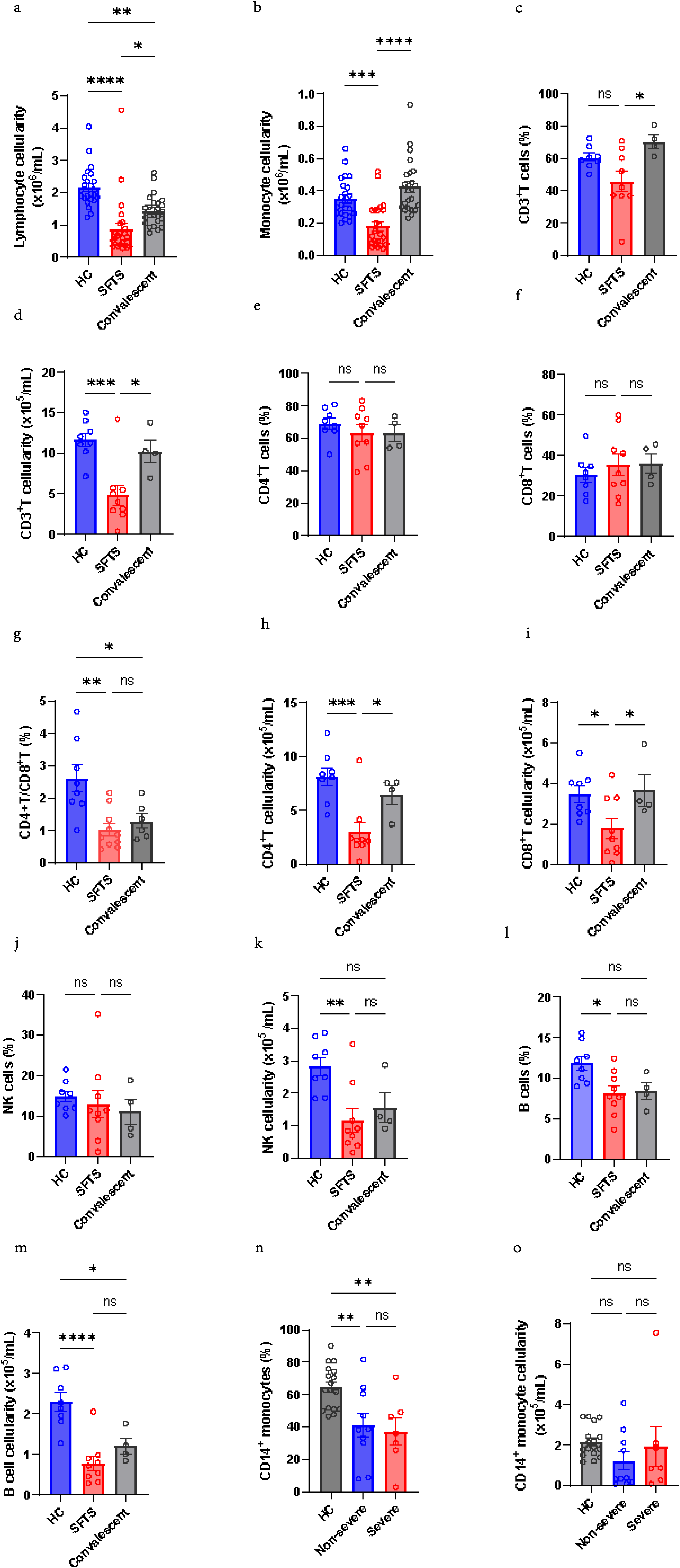
Altered PBMC composition in patients with SFTS. (a-m) Percentages and absolute values of PBMCs examined by flow cytometry of each sample from the HC group, SFTS group and Convalescent group. (n-o) Percentages and absolute values of CD14^+^ monocytes examined by flow cytometry of each sample from the HC group and two SFTS groups (Non-severe, Severe). Continuous variables are shown as mean and standard deviation or median and interquartile.

Corroborating previous findings, monocytes in SFTS patients experienced a substantial decrease in PBMCs and subsequently restored during the convalescent stage (Fig. 3b). Likewise, both the ratio and cell count of CD14^+^ monocytes in SFTS patients showed a decline (Fig. 3n,o).

### Identification of monocyte heterogeneity

Monocytes play a critical role in controlling SFTSV infection in patients. To further describe the heterogeneity of human blood monocytes at the single-cell level and provide additional markers for more precise subset classification, we performed scRNA-seq on sorted monocytes from PBMCs of seven SFTS patients. Additionally, we used scRNA-seq data from three normal monocyte samples sourced from public databases for comparative analysis. We analyzed 59,901 circulating monocytes and identified nine distinct populations based on transcriptome profiles (Fig. 4a,b). By examining the expression patterns of CD14 and FCGR3A/CD16, we classified five subsets as CD16^+^ and four as CD14^+^ monocytes (Fig. 4c).

**Fig. 4.**
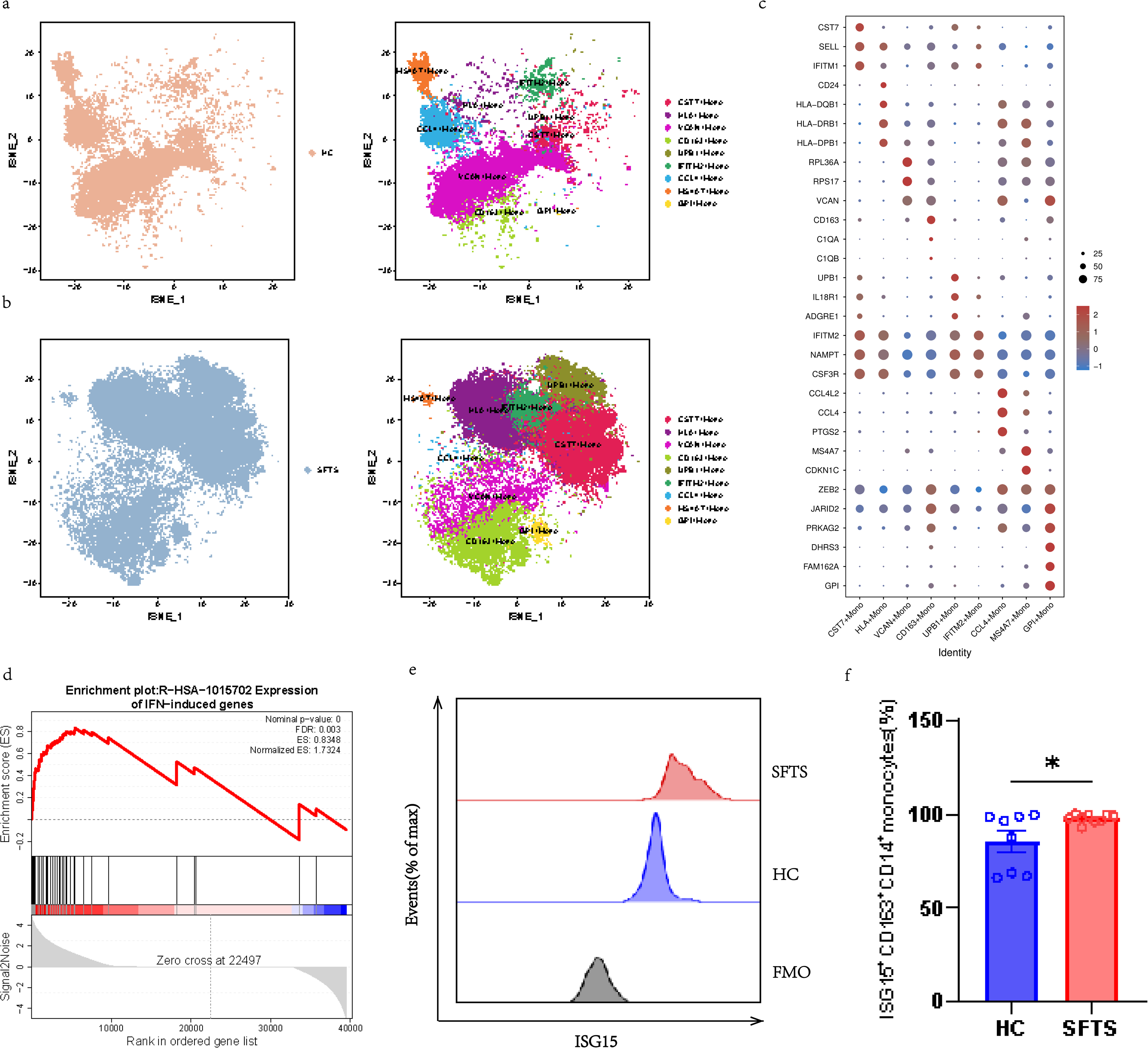
Identification of monocyte subsets among SFTS patients via scRNA-seq transcriptome analyses. (a-b) UMAP plots showing cell clustering results of the monocyte compartment from sorted monocytes from blood samples of SFTS patients (a, n = 7) and HCs (b, n = 3). (c) Dot plot shows the expression percentages and the expression levels of feature genes of different clusters of monocytes. (d) GSEA analysis show IFN-induced genes enrichment in SFTS group. (e-f) The intracellular expression of ISG15 on CD163^+^CD14^+^ monocytes of the HC group and SFTS patients. Student’s t-test was performed, according to the data distribution. \**p*< 0.05.

We then compared the functional phenotypes of all nine subtypes by assessing the highly expressed genes (Fig. 4c). Specifically, CST7-high monocytes (CST7+Mono) overexpressed CST7 (encoding cystatin F), which is upregulated in a subset of disease-associated microglia in Alzheimer’s disease models [23]. While the expression profile of IFITM2-high monocytes (IFITM2+Mono) was similar to that of CST7+Mono, IFITM2+Mono cells were enriched in responses to oxidative stress, expressing redox-related genes (SOD2 and NAMPT) important for antiviral signaling regulation [24]. UPB1-high monocytes (UPB1+Mono) involved inflammation and immunity genes (UPB1 and C15orf48) [25]. MS4A7-high monocytes (MS4A7+Mono) were nonclassical monocytes; besides confirming FCGR3A expression, we also identified markers such as MS4A7, CDKN1C, CSF1R, HES4, RHOC, and MTSS1, indicating enrichment in negative regulators of cell proliferation, genes associated to the Wnt signaling pathway, Notch signaling pathway and type I interferon antiviral response [26–30]. CCL4-high monocytes (CCL4+Mono) showed a different gene combination, including chemokine ligands, nod-like receptors, and interleukins (e.g., CCL4, CCL3, CXCL2, NLRP3, and IL1B) related to viral infection, the inflammatory process, and pyroptotic cell death [31–33]. Human leukocyte antigen (HLA)-high monocytes HLA+Mono) were enriched for genes in antigen presentation pathways, such as HLA class II alleles (HLA-DQB1, HLA-DRB1, HLA-DPB1, and CD24). CD163-high monocytes (CD163+Mono) showed preferential expression of macrophage markers (CD163) (Fig. 4c) and IFN-inducible genes (ISG15, IFI27, IFI6, and MX1) (Fig. 4d). Previous studies have shown that interferon-stimulated gene 15 (ISG15), a member of the ubiquitin family, strongly upregulated during viral infections and exerts pro-viral or antiviral actions [34]. However, there is no report on the role of ISG15 in SFTS. Hence, we further explored the expression of ISG15 at the protein-level in CD163+Monos using flow cytometry. Our results demonstrated significantly higher levels of ISG15 in SFTS patients than in healthy controls (Fig. 4e,f). Among the three subsets with lower CD14 expression, VCAN-high monocytes (VCAN+Mono) overexpressed VCAN gene, and GPI-high monocytes (GPI+Mono) showed overexpression of GPI gene (Fig. 4c).

We then compared the differences in monocyte composition between the SFTS and HC groups (Fig. 5a,b). Comparing the proportions of monocyte subtypes in the SFTS and HC groups, we noted a decrease in CD14^+^ monocytes among SFTS patients (Fig. 5c,e), which consistent with the the results from flow cytometry (Fig. 3n,o). Notably, there was a significant increase in CD163+Mono cells (Fig. 5d,f) and a decrease in CCL4+Mono cells (Fig. 5d) in the SFTS group. Using flow cytometry, we validated that the CD163^+^CD14^+^ monocytes (CD163+Mono cluster in scRNA-seq data) was increased in SFTS patients (Fig. 5g). We then investigated HLA-DR expression, known as an activation marker, given its reduced levels in acute severe COVID-19 and sepsis patients as part of an immune-protective mechanism [35–36]. Compared to non-severe hospitalized SFTS patients, severe SFTS cases demonstrated reduced HLA-DR expression on circulating CD14^+^ monocytes and CD163^+^CD14^+^ monocytes (Fig. 5h-k), indicating a dysfunctional immune response.

**Fig. 5.**
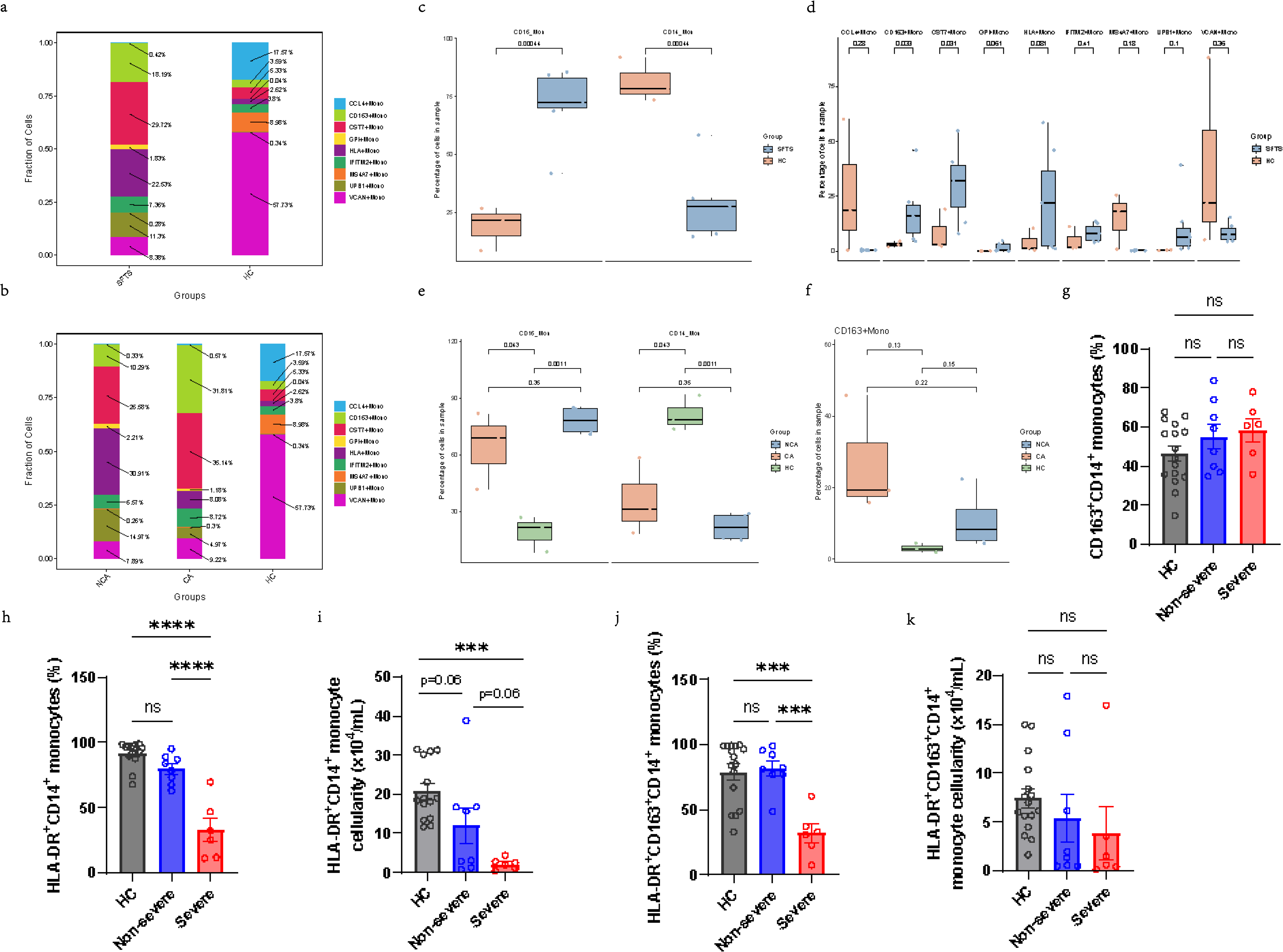
Identification of monocyte subsets among SFTS patients via scRNA-seq transcriptome analyses. (a) Bar plots showing cluster distribution of the monocyte subsets in SFTS patients and HCs (colors are coded for the cell subpopulations identified in this study). (b) Proportional bar charts of cell subpopulations in SFTS patients (NCA and CA) and healthy controls. (c-d) Graph showing percentages of monocyte subsets in SFTS patients and HCs. (e-f) Boxplot comparing the differences in cell proportions between SFTS patients (NCA and CA) and HCs in the sorted monocyte scRNA-seq data. (g) The proportions of CD163^+^CD14^+^ monocytes in peripheral monocytes from SFTS patients and HCs. (h-k) HLA-DR expression on CD14^+^ monocytes and CD163^+^CD14^+^ monocytes in healthy, non-severe and severe groups based on flow cytometric analyses of PBMCs. CA, critical group at the acute phase; NCA, non-critical group at the acute phase. Data are presented as the mean±SEM. The p-values were calculated with nonparametric one-way ANOVA (Kruskal-Wallis test) followed by Tukey’s multiple comparison test. \**P* < 0.05; \*\**P* < 0.01; \*\*\**P* < 0.001; \*\*\*\**P* < 0.0001. ns, not significant.

Overall, analysis of scRNA-seq data for sorted monocytes demonstrated the heterogeneity and functional variation among circulating monocytes, and significantly increased CD163^+^ monocytes in patients with SFTS.

### Differentiation trajectories toward CD163+Mono cluster

To investigate the developmental relationships among monocyte subtypes, we established the pseudotemporal order and reconstructed the differentiation trajectory using Monocle 2 [17]. The pseudotime trajectory originated from HLA+Mono (state 0) and divided into two branches: state 1, characterized by CCL4+Mono, and state 2, identified as CD163+Mono (Fig. 5a,b). Notably, a marked imbalance in distribution was observed between the two conditions across these states, with SFTS monocytes predominating at the end of state 2, while more HC monocytes accumulated at the conclusion of state 1 (Fig. 6a,b). This trend corresponded with the elevated levels of CD163+Monos and reduced levels of CCL4+Monos signatures in SFTS (Fig. 5d).

**Fig. 6.**
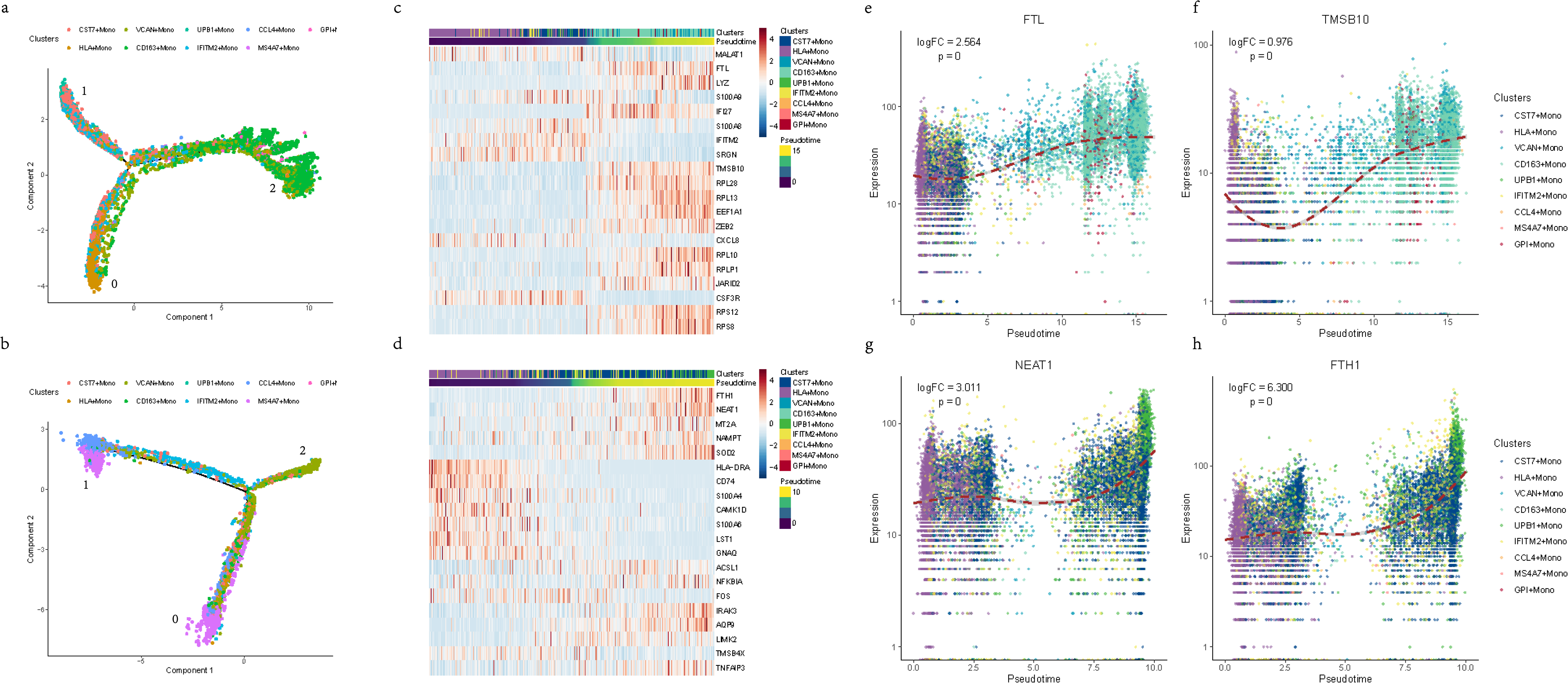
Trajectory inference of monocyte differentiation. (a-b) Visualization of monocyte trajectories from SFTS group (a) and HC group (b) using Monocle 2, colored by monocyte subtype and pseudotime. Each dot represents a cell. (c-d) Heatmap showing the gene expression (normalized log count, shown by color) of correlated genes along state 1 (d) and state 2 (c), annotated by the corresponding pseudotime and cell type. (e-h) Expression of known TFs driving macrophage development along state 2 (e,f) and state 1 (g,h), colored by monocyte subtypes. Dashed line shows the smoothed fit of TF’s expression along state. Labeled are *P* value and log fold-change (logFC) of the TF along state. *P*.adj, adjusted *P* value.

Subsequently, we investigated the associated gene expression profiles and pathways potential these two differentiation trajectories. State 2 exhibited variations in genes related to innate immunity (FTL, IFI27, and TMSB10) (Fig. 6c). Conversely, there was a progressive upregulation of genes encoding iron metabolism-related proteins (FTH1, NEAT1, and IRAK3) along state 1 (Fig. 6d). Examining the expression of key transcription factors (TFs) plotted along the pseudotime axis, we observed that protease genes upregulated during state 2 differentiation included FTL and TMSB10 (Fig. 6e,f) [37]. Studies have indicated that FTL induces M2 macrophage polarization and that TMSB10 promotes M2 conversion and proliferation through the PI3K/Akt pathway [38]. On the other hand, TFs involved in the monocyte-to-macrophage transition (NEAT1 and FTH1) displayed a gradual increase in expression in state 1 (Fig. 6g,h). NEAT1 has been found to regulate hantaan virus (HTNV)-induced M1 polarization, inhibiting viral propagation and cell-to-cell spread. Additionally, NEAT1 expression in monocytes has been negatively correlated with progression in hemorrhagic fever with renal syndrome (HFRS) [39]. FTH1 serves as a marker gene for M1 macrophages, participating in macrophage polarization by regulating iron metabolism, with higher expression in M1 compared to M2 macrophages [40–41]. In summary, genes associated with M2 characteristics drive monocytes towards differentiation into CD163+Mono cells during SFTSV infection and may be related to disease severity.

### Activated IL-6/JAK/STAT3 signaling of CD163^+^ monocytes in SFTS

To understand the mechanisms driving the expansion of CD163+Mono cells in SFTS, we initially conducted an analysis of Differentially Expressed Genes (DEGs) in CD163^+^CD14^+^ monocytes between SFTS patients and HCs, identifying 4262 up-regulated genes and 1303 down-regulated genes in SFTS (Fig. 7A). Gene Ontology (GO) analysis indicated that genes associated with the inflammatory response, IL-6/JAK/STAT3 signaling, and interferon α/γ response were commonly up-regulated during infection, while genes involved in oxidative phosphorylation, Wnt/β-catenin signaling, and TNF-α signaling via NFκB were significantly down-regulated during infection (Fig. S1).

**Fig. 7.**
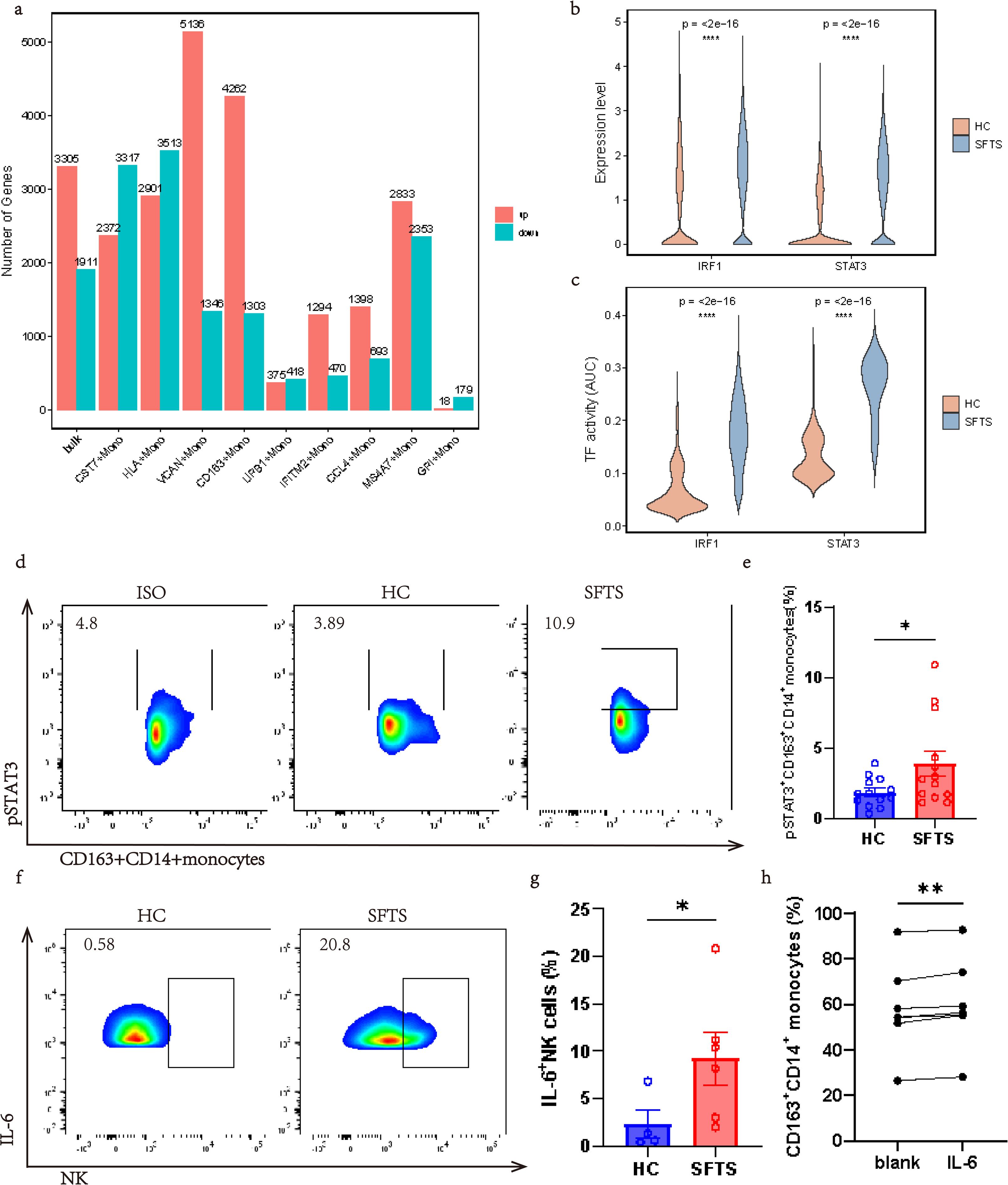
Activated IL-6/JAK/STAT3 signaling of CD163^+^ monocytes in SFTS. (a) Distribution of the number of genes detected in each subset in scRNA-seq data of PBMCs. (b) Violin plot comparing the difference in the area under the curve (AUC, from the SCENIC algorithm) for the TFs between SFTS patients (blue) and HCs (red). (c) Violin plot showing the expression of predicted TFs (IRF1 and STAT3) in CD163+Mono of SFTS patients and HCs. (d-e) Escalating pSTAT3 levels of CD163^+^CD14^+^ monocytes in SFTS. Representative flow cytometry data (d) and averaged percentages of (e) CD163^+^CD14^+^ monocytes. (f-g) Representative flow cytometry data (f) and averaged percentages (g) of IL-6^+^ NK cells of HC and SFTS patients. (h) Graph displaying the proportions of CD163^+^CD14^+^ monocytes of IL-6 stimulation.

Next, we used SCENIC tools [19] to predict the Transcription Factors (TFs) regulating these DEGs and observed a notable enrichment of TFs that regulating IL-6/JAK/STAT3 signaling (Fig. S2). Among the up-regulated DEGs, Interferon-Stimulated Genes (ISGs) like IRF1 playing important roles in the IFN response post-monocyte exposure (Fig. 7b,c). STAT3, a key transcription factor in the JAK-STAT pathway related to cytokine receptor signaling [42], showed elevated expression in CD163^+^CD14^+^ monocytes from SFTS patients (Fig. 7b,c). Flow cytometry confirmed a significant increase in STAT3 phosphorylation within CD163^+^CD14^+^ monocytes from SFTS patients (Fig. 7d,e).

The enriched IL-6/JAK/STAT3 signaling in SFTS CD163^+^CD14^+^ monocytes prompted an investigation into whether IL-6 could stimulate these cells, revealing a significant increase in the proportion of CD163^+^CD14^+^ monocytes following IL-6 treatment (Fig. 7h).

Furthermore, elevated IL-6 levels were detected in serum from SFTS patients (Fig. 1b). Analysis of IL-6-producing cells showed a significant increase in IL-6-positive NK cells (Fig. 7f,g). Exploring the regulatory connections among cell clusters, we conducted cell-cell communication analysis of available single-cell transcriptome data to infer potential interactions between CD163+Mono cells and TBNK cells. Notably, incoming interactions from TBNK cells to CD163+Mono cells were predominantly increased in NK cells from SFTS patients, consistent with heightened IL-6 production by NK cells. These analyses illustrated frequent crosstalk between CD163+Mono cells and NK cells in SFTS patients, indicating potential anti-inflammatory characteristics in SFTS-associated interactions, such as THBS1-CD47 [43–44], LGALS9-CD45 [45], ADGRE5-CD55 [46], and ANXA1-FPR1 [47] (Fig. S4).

### Tofacitinib inhibits IL-6-induced monocyte M2 polarization in patients with SFTS

Moreover, we investigated the response of CD163^+^CD14^+^ monocytes in SFTS to JAK inhibition. The JAK-STAT pathway signaling represent a classical pathway through which IL-6 activates STAT3 TF [48]. Given the evident activation of STAT3 signaling in CD163^+^ monocytes (Fig. 7b-e), our subsequent inquiry focused on determining the impact of tofacitinib (a JAK inhibitor) on CD163^+^CD14^+^ monocytes. Flow cytometry analysis indicated an increase in the proportion of CD163^+^ monocytes in monocyte cultures treated with IL-6 compared to those treated with PBS (Fig. S5). Similarly, flow cytometry demonstrated a significant decrease in proportions following treatment of monocytes with tofacitinib and IL-6 (Fig. S5). These findings show that tofacitinib treatment effectively inhibits the differentiation of monocytes toward the M2 phenotype.

### CD163+Mono cells are the major cell types that infected by SFTSV

By aligning our single-cell RNA sequencing (scRNA-seq) findings with a custom genome containing human and SFTSV sequences, we pinpointed cells containing viral mRNA. Our study revealed the presence of SFTSV transcripts exclusively in the critical patient group, not in the non-critical group. Among approximately 55 viral transcript-containing cells in the critical cohort, surprisingly, 26 were identified as CD163+Mono cells (Fig. 8a,b). Additionally, the viral load in serum samples from critically ill patients significantly surpassed that of non-critical cases (Fig. 8c), indicating a strong correlation between SFTSV replication levels and disease severity, further supported by trajectory analysis.

**Fig. 8.**
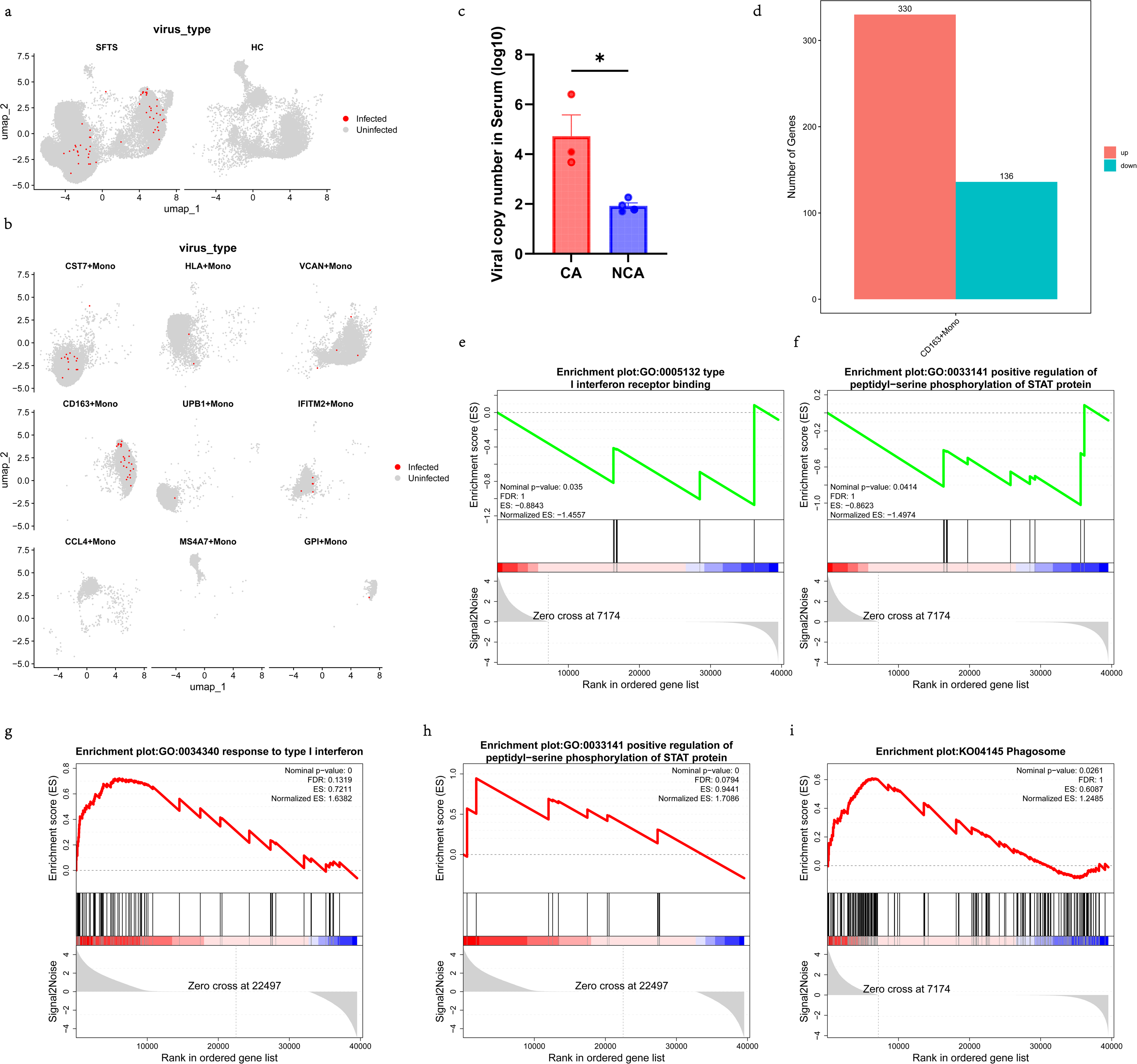
CD163+Mono cells are the primary viral reservoir in severe cases of SFTSV infection. (a) t-SNE plots of two groups. Cells with SFTSV transcripts are colored red, while cells without SFTSV transcripts are colored gray. (b) t-SNE plots of monocyte subtypes with SFTSV^+^ cells (red) and SFTSV^-^ cells (gray) in SFTSV infections. (c) Quantification of SFTSV copy numbers in patient sera using RT-PCR. The viral copy numbers in sera were higher in the CA group than in the NCA group. CA, critical group at the acute phase; NCA, non-critical group at the acute phase. (d) A bar plot shows that 330 genes were upregulated and 136 genes were downregulated in infected CD163+Mono cells compared to uninfected CD163+Mono cells. (e-f, i) GSEA analysis show differential expression pathways enrichment in SFTSV-infected CD163+Mono cells versus non-infected CD163+Mono cells in SFTSV infections. (g-h) GSVA plot showing differentially expressed genes in the SFTS group versus HC group in CD163+Mono cluster.

Assessing the impact of SFTSV infection on monocyte-driven immune responses, we scrutinized gene expression changes in infected CD163+Mono cells compared to uninfected counterparts within the SFTS group. Notably, a bar plot demonstrated 330 upregulated genes and 136 downregulated genes in infected CD163+Mono cells versus uninfected ones (Fig. 8d). Gene set enrichment analysis (GSEA) highlighted a downregulation of the IFN signaling pathway and the JAK-STAT signaling pathway in infected CD163+Mono cells relative to their uninfected counterparts in the SFTS group (Fig. 8e,f). Conversely, there was an upregulation of Phagosome-related genes in infected CD163+Mono cells (Fig. 8i), highlighting their essential role in pathogen clearance within the innate immune system.

## Discussion

In the present study, we demonstrated immunosuppression of essential innate immune cells in severe SFTS patients, expression of MHC class II (human leukocyte antigen-DR [HLA-DR]) on circulating CD14^+^ monocytes was reduced, and this was not observed in hospitalized SFTS patients without critical illness. Immunoparalysis, noted as a persistent anti-inflammatory response to severe insults like trauma or sepsis, amplifies the vulnerability to opportunistic infections and higher morbidity and mortality risks. This downregulation of HLA-DR expression is not restricted to infectious diseases and is commonly observed in critically ill patients post-trauma, burn injuries, major surgeries, or during pancreatitis [49–50]. A study has shown that monitoring changes in HLA-DR^+^ monocytes can distinguish survivors from non-survivors among sepsis patients [51]. Lukaszewicz et al., present a study that monitoring immune functions through mHLA-DR in ICU patients could be useful to identify a high risk of secondary infection [52]. Therefore, low HLA-DR on monocytes may be more than a prognostic marker of disease severity. In several studies, monocytic HLA-DR expression has been utilized as an indicator of the efficacy of immunostimulatory therapies [53–55].

The observed immunosuppression associated with decreased HLA-DR expression could contribute to increased viral replication or disease severity progression. Previous data from critical patients with bacterial septic shock suggests higher rates of secondary infections in patients with consistent HLA-DR downregulation, characteristic of injury-related immunosuppression [56]. Given the impact of secondary infections on mortality in critical SFTS patients [57–58], further research should be investigated in the potential relation between HLA-DR downregulation and secondary infection rates.

Utilizing high-resolution single-cell technologies, we redefined monocytes into nine subtypes, exploring the pathogenic role of these subtypes in SFTS. Our experiments showed the expansion of CD163^+^ monocytes in SFTS patients, correlating with disease severity. Trajectory analysis revealed the preferential differentiation of SFTS monocytes into CD163^+^ monocytes. Moreover, our study identified CD163^+^ monocytes as main targets in critical patients through scRNA-seq analysis of SFTSV infectivity in peripheral blood samples, supporting the conclusion that macrophages shift towards an M2 phenotype under SFTSV infection [59]. Most monocyte-tropic viral infections, like HIV, RSV, SARS, and IAV, alter macrophage polarization, leading to immunosuppression and potential immunopathology. This can result in viral persistence and co-infections with pathogens from different groups [60]. Understanding these phenotypic and functional changes in monocytes may provide insights into the pathogenesis of SFTS.

The mechanisms by which IL-6 signaling mediates immune disturbances in SFTS remain unclear, despite significantly elevated IL-6 levels in SFTS patients. Notably, IL-6 can exhibit either pro-inflammatory [61] or anti-inflammatory [62] effects depending on the local immune microenvironment. Activation of STAT3 by IL-6 triggers downstream effects, thereby increasing the transcription and expression of various target genes [63]. Our study revealed that CD163^+^ monocytes respond to IL-6 activation in SFTS patients. As a critical transcriptional modulator of IL-6, STAT3 was also prominently activated in CD163^+^ monocytes, similar to that in active macrophages [64] in other diseases. Additionally, treatment with IL-6 induced an increase in CD163^+^ monocytes, consistent with previous findings [65].

The high mortality rate in SFTS generally associated with lymphopenia and a severe inflammatory response triggered by unregulated cytokine release. These mediators are regulated transcriptionally via the JAK/STAT signaling pathways, which can be modulated by small molecules. Targeting the JAK-STAT pathway, whether alone or in combination with antiviral drugs, has shown promising efficacy in pre-clinical studies and phase 1 trials. It has been reported that baricitinib and ruxolitinib improved clinical outcomes for COVID-19 patients by reshaping the immune response and mitigating immunosuppression in myeloid cells [66–68]. A study also indicate the efficacy and safety of ruxolitinib in treating cytokine storms in viral infections, with potential benefits for survival rates and reduced ICU hospitalizations in severe SFTS cases [69]. Our results suggest that tofacitinib stimulation dramatically inhibited the polarization of monocytes toward the M2 phenotype. Clinical trials incorporating JAK inhibitors, possibly in combination with antiviral drugs, are essential for determining optimal dosing, efficacy, and tolerability in SFTS patients.

## Conclusion

In conclusion, this study provided detailed single-cell profiles of monocytes in SFTS patients. By comparing these profiles with those of healthy individuals, we identified changes in monocyte subgroup distributions in SFTS patients. Additionally, SFTSV activates the JAK/STAT3 pathway, leading to an upregulation of CD163^+^ monocytes proportion that convert monocytes to the M2 phenotype. Investigating and validating JAK inhibitors is crucial for developing effective strategies against SFTS and other emerging tick-borne hemorrhagic fevers. Given the emergence of bunyaviridae viruses like Yezo virus (YEZV) and Wetland virus (WELV) worldwide [70–71], understanding the defined features of SFTS may enhance our comprehension of immunopathogenesis for other bunyaviridae viruses.

## Acknowledgements

YZ was supported by the China Scholarship Council (grant no. 202408340092).

## Author Contributions

Experiments in this study were conceived by YZ. Samples were collected by YZ, QY YJ and YL. Experiments were performed by YZ, and MG. Data analysis was performed by YZ. The manuscript was written by YZ, and edited by YX. All authors read and approved the final manuscript. YZ and MG contributed equally to this work.

## Data availability

Data will be made available on request. Publicly available datasets used in this manuscript are available from GSE287827.

---

**Fig. S1.** Heatmap showing DEGs in CD163+Mono between the SFTS and HC groups.

**Fig. S2.** Heatmap of the AUC scores of expression regulation by transcription factors in CD163+Mono. AUC, area under the curve.

**Fig. S3.** Cell-cell communication analysis reveals robust NK cell-monocyte cross-talk in SFTS. A scRNA-seq dataset from Li et al. [58] was analyzed using the CellPhoneDB.

**Fig. S4.** Cell-cell interaction analysis between CD163+Mono clusters and other immunecell clusters in SFTS group.

**Fig. S5.** IL-6-stimulated CD163^+^CD14^+^ monocytes were decreased after tofacitinib treatment, detected by flow cytometry. The Friedman test was applied. \*\**P* < 0.01.

