## Supplementary figures and images for "Analysis of SFTS pathogenesis based on single-cell RNA sequencing of monocytes"

### FigureS1.tif

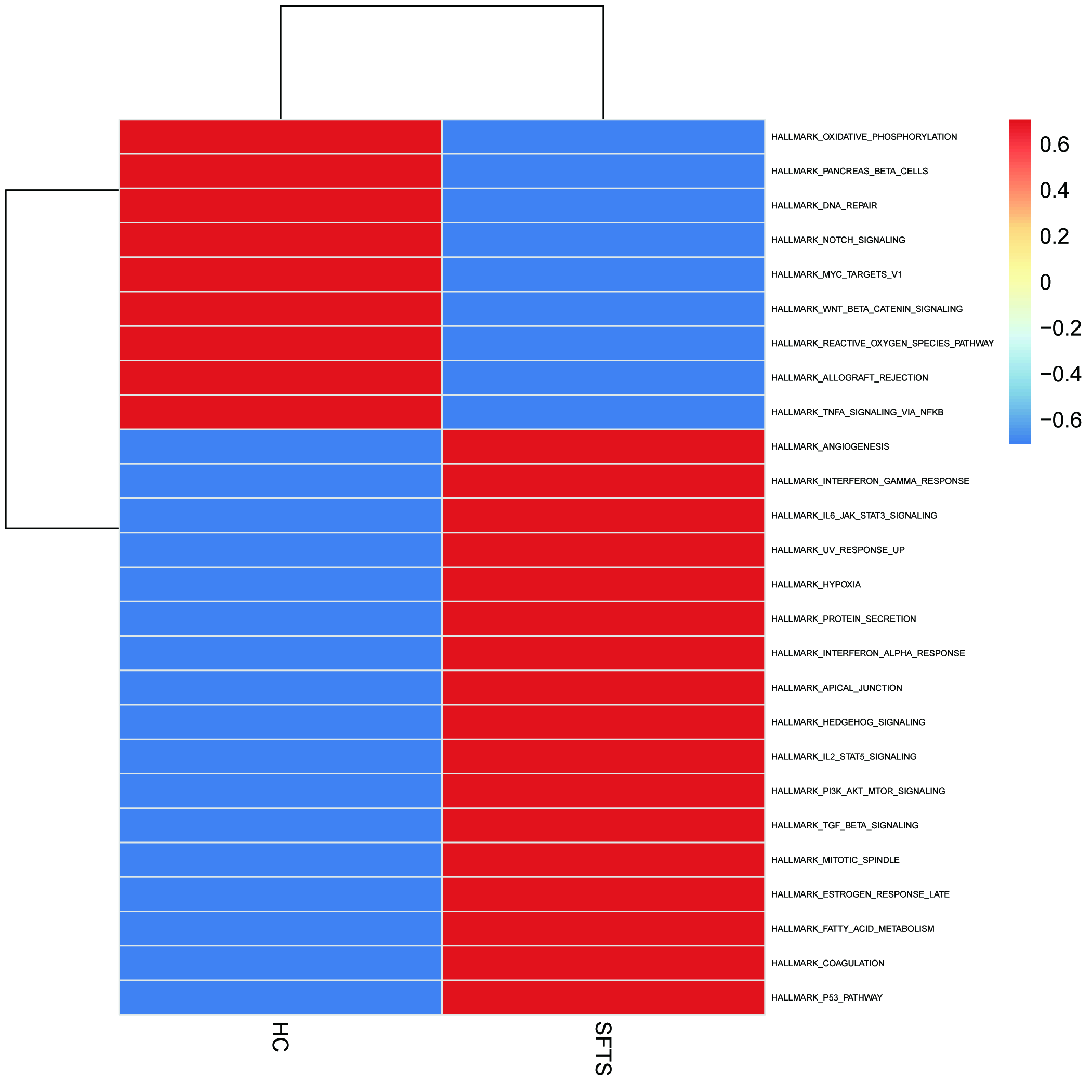

### FigureS2.tif

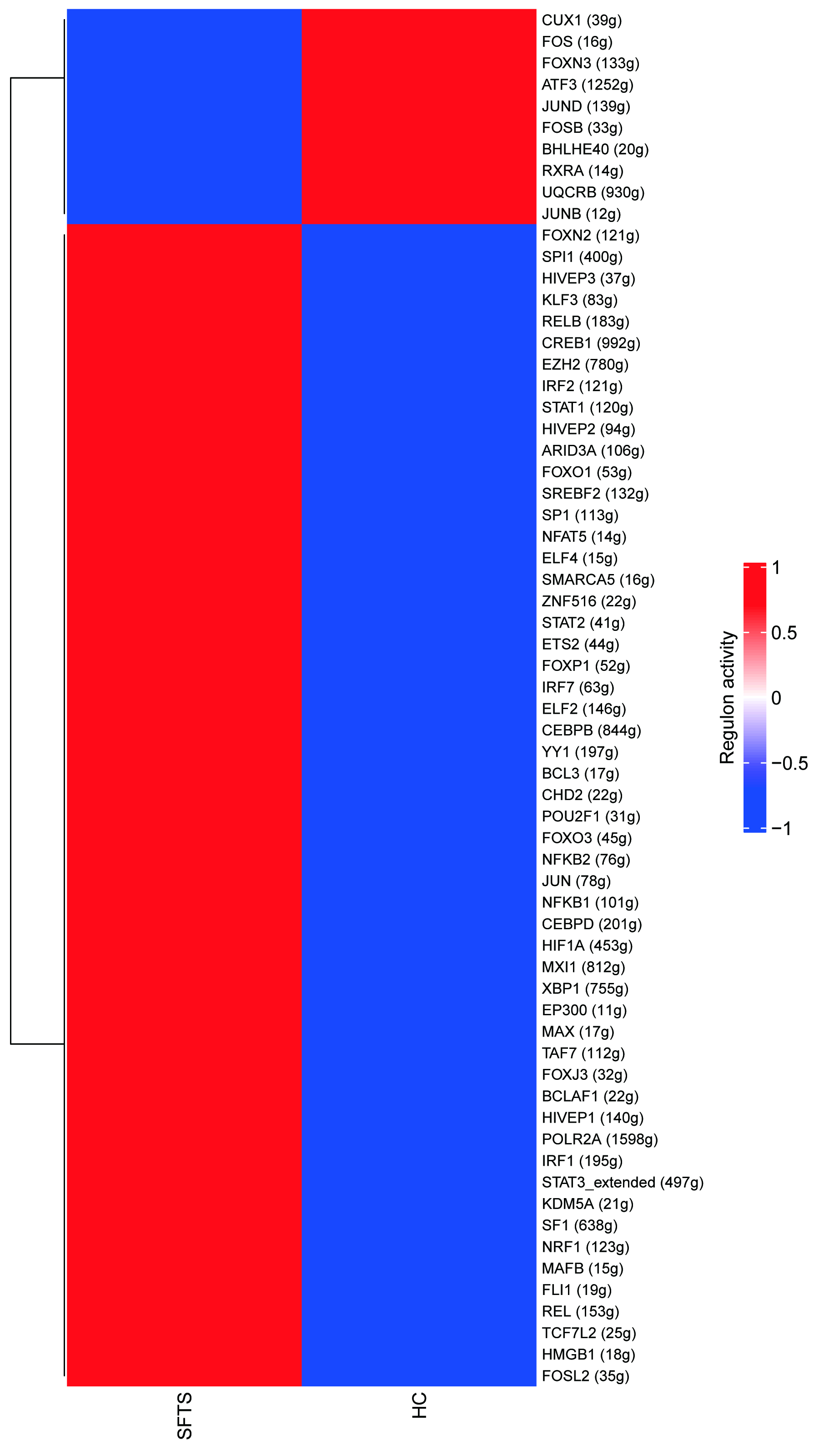

### FigureS3.tif

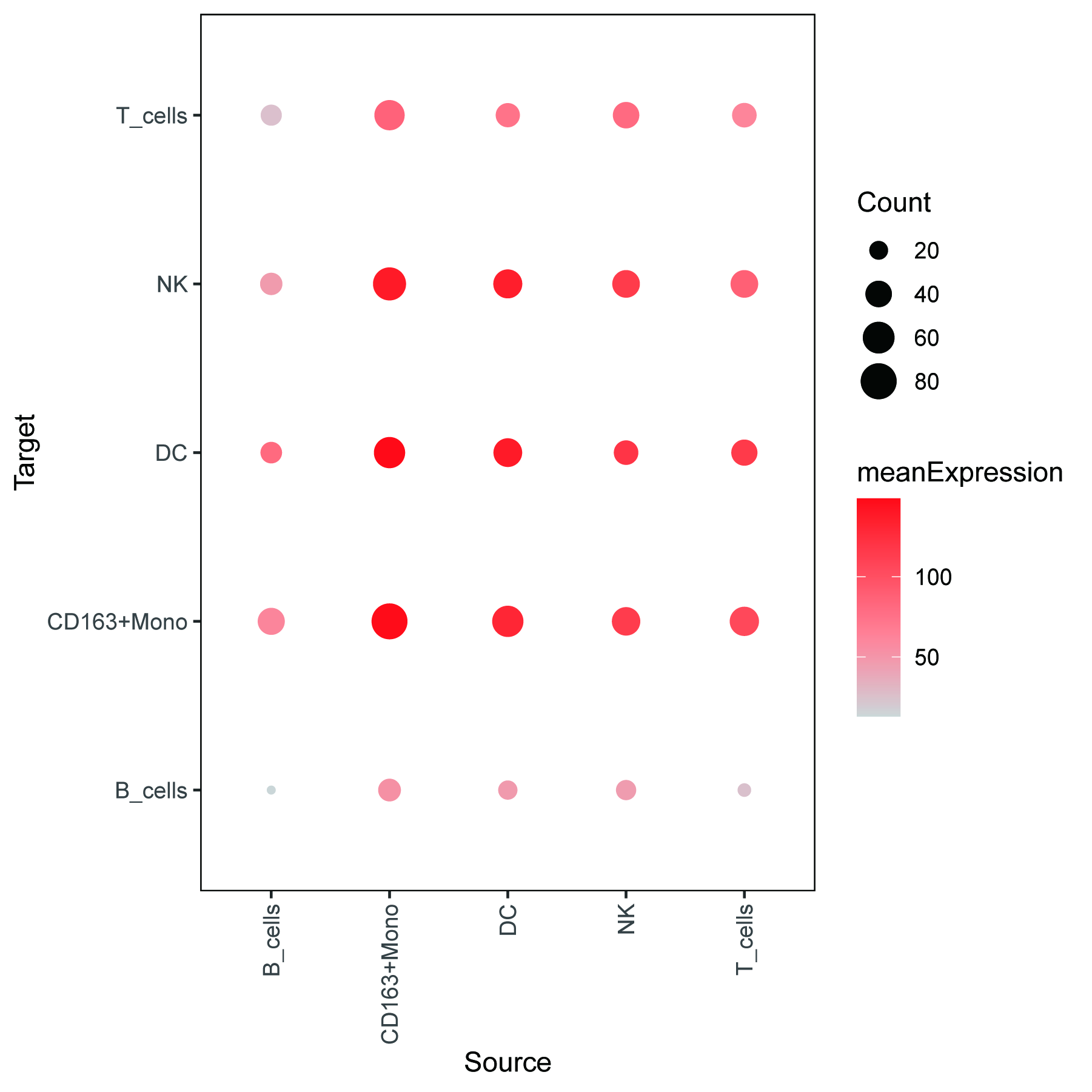

### FigureS5.tif

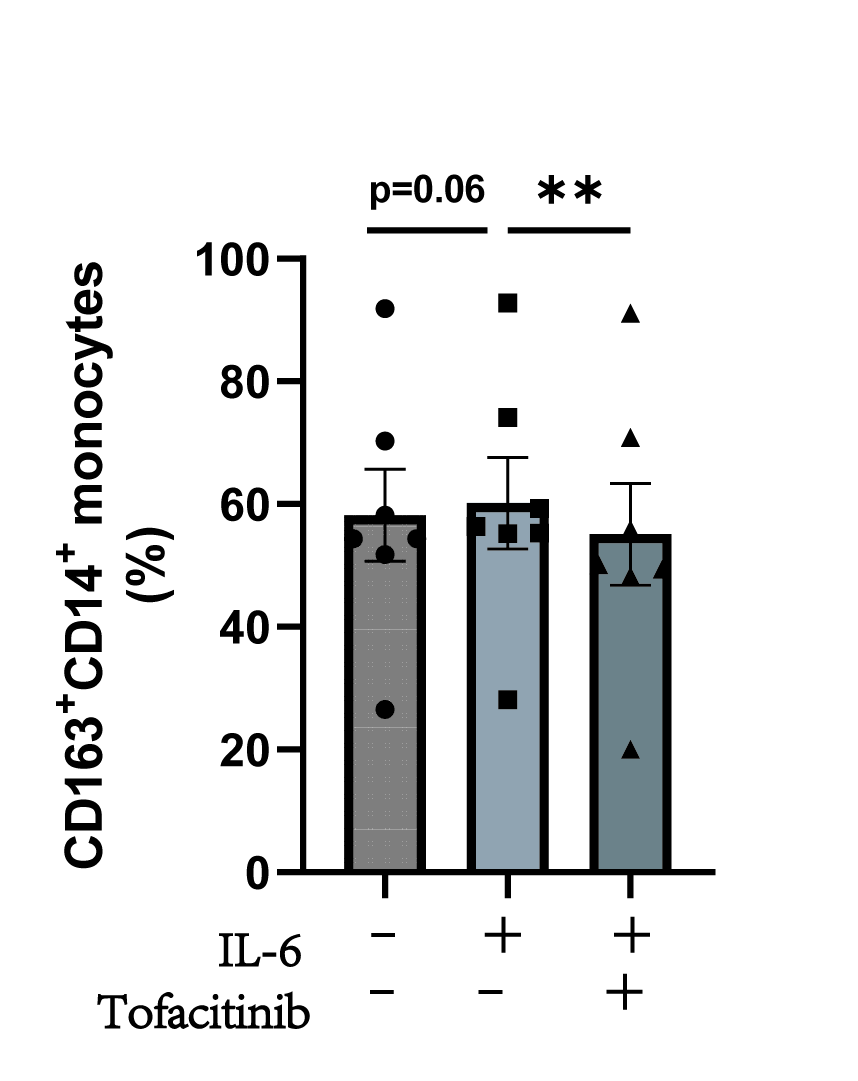
